# Cortical Coding of Gustatory and Thermal Signals in Active Licking Mice

**DOI:** 10.1101/2024.04.27.591293

**Authors:** Audrey N. Nash, Morgan Shakeshaft, Cecilia G. Bouaichi, Katherine E. Odegaard, Tom Needham, Martin Bauer, Richard Bertram, Roberto Vincis

## Abstract

Eating behaviors are influenced by the integration of gustatory, olfactory, and somatosensory signals, which all contribute to the perception of flavor. Although extensive research has explored the neural correlates of taste in the gustatory cortex (GC), less is known about its role in encoding thermal information. This study investigates the encoding of oral thermal and chemosensory signals by GC neurons compared to the oral somatosensory cortex. In this study, we recorded the spiking activity of more than 900 GC neurons and 500 neurons from the oral somatosensory cortex in mice allowed to freely lick small drops of gustatory stimuli or deionized water at varying non-nociceptive temperatures. We then developed and used a Bayesian-based analysis technique to assess neural classification scores based on spike rate and phase timing within the lick cycle. Our results indicate that GC neurons rely predominantly on rate information, although phase information is needed to achieve maximum accuracy, to effectively encode both chemosensory and thermosensory signals. GC neurons can effectively differentiate between thermal stimuli, excelling in distinguishing both large contrasts (14°C vs. 36°C) and, although less effectively, more subtle temperature differences. Finally, a direct comparison of the decoding accuracy of thermosensory signals between the two cortices reveals that while the somatosensory cortex showed higher overall accuracy, the GC still encodes significant thermosensory information. These findings highlight the GC’s dual role in processing taste and temperature, emphasizing the importance of considering temperature in future studies of taste processing.

**Key Points:**

- Flavor perception relies on gustatory, olfactory, and somatosensory integration, with the gustatory cortex (GC) central to taste processing.
- GC neurons also respond to temperature, but the specifics of how the GC processes taste and oral thermal stimuli remain unclear.
- The focus of this study is on the role of GC neurons in the encoding of oral thermal information, particularly compared to the coding functions of the oral somatosensory cortex.
- We found that while the somatosensory cortex shows a higher classification accuracy for distinguishing water temperature, the GC still encodes a substantial amount of thermosensory information.
- These results emphasize the importance of including temperature as a key factor in future studies of cortical taste coding.

## Introduction

Eating behaviors are influenced by the initial sensation and the reward experienced when eating food and beverages. This sensation results from the integration of intraoral gustatory, olfactory (retronasal), and somatosensory cues that all contribute to the percept that we know as flavor (Samuelsen and Vincis, 2021; Spence, 2015; Small, 2012; Lemon, 2021). Numerous electrophysiological studies in behaving rodents have described the neural correlates of one of these sensory components, taste. Using fluid stimuli at room temperature, these investigations revealed that gustatory information undergoes neural computations within interconnected brain regions, including the gustatory cortex (GC), the primary cortical region responsible for the processing of taste (Vincis and Fontanini, 2019; Spector and Travers, 2005). GC neurons have shown time-varying patterns of activity in response to chemosensory qualities and the hedonic value of gustatory stimuli, which play a role in guiding taste-related decisions (Katz et al., 2001; Stapleton et al., 2006; Dikecligil et al., 2020; Jezzini et al., 2013; Levitan et al., 2019). Furthermore, the ability of GC neurons to distinguish between tastes is improved when the rate and spike-time are interpreted relative to the timing of the licks, indicating that the lick cycle is a key factor in taste processing (Neese et al., 2022).

Several studies indicate that GC neurons also respond to non-gustatory components of oral stimuli (Vincis and Fontanini, 2016; Samuelsen and Fontanini, 2017; Rudenga et al., 2010; Samuelsen and Vincis, 2021), including temperature, a prominent feature of the sensory properties of food and beverages. Studies in humans and primates (Cerf-Ducastel et al., 2001; Verhagen et al., 2004), as well as in anesthetized rats (Yamamoto et al., 1981; Kosar et al., 1986) and investigations by our laboratory in awake mice (Bouaichi et al., 2023), suggest that changes in oral temperature, even in the absence of classical taste qualities, modulate activity in GC neurons. In particular, our recent findings (Bouaichi et al., 2023) revealed that more than half of the GC neurons that encode chemosensory taste stimuli can also discriminate thermal information. Although these data suggest that the GC is a potential key brain region for integrating chemosensory and thermosensory inputs from the oral cavity, they do not provide a detailed analysis of its neural responses. Consequently, different aspects of the GC’s capability to process taste and oral thermal stimuli are poorly characterized. For example, it is still unknown what coding strategy is used by GC neurons in the encoding of oral thermal information, particularly compared to the thermosensory functions of the oral somatosensory cortex (S), which represents the sensory input of the tongue and intraoral region (Accolla et al., 2007; Nakamura et al., 2015; Clemens et al., 2018; Samuelsen and Vincis, 2021)

This study aims to evaluate the role of GC neurons in the encoding of oral thermal information and their ability to process chemosensory taste signals at room temperature, particularly compared to the thermosensory and chemosensory coding functions of the oral somatosensory cortex (S), which represents the sensory input of the tongue and intraoral region (Accolla et al., 2007; Nakamura et al., 2015; Clemens et al., 2018; Samuelsen and Vincis, 2021). To this end, we collected recordings of spiking activity from more than 900 GC neurons and 500 neurons in the somatosensory cortex in mice allowed to freely lick to receive a small drop (3 *µ*l) of one of four liquid gustatory stimuli (sucrose, NaCl, citric acid, and quinine) at room temperature or deionized water at one of three different non-nociceptive temperatures (14°, 25° and 36°C). Our previous study (Bouaichi et al., 2023) indicated that the responses of GC neurons to different temperatures of deionized water and artificial saliva - a stimulus often used as a neutral control in taste research - were highly similar, supporting the use of deionized water at different temperatures as a thermal stimulus in the absence of overt chemosensory taste information. We then applied a Bayesian analysis to compute classification scores for spike trains, considering both the rate and the phase (timing of spikes within the lick cycle) in response to various oral stimuli. Our results indicate that GC neurons primarily rely on rate information, with phase providing complementary input, to effectively encode chemosensory and thermosensory signals from the oral cavity. In addition, a qualitative evaluation of our decoding analysis showed that many GC neurons are able to distinguish different water temperatures. Finally, a direct comparison of the decoding accuracy of thermosensory and chemosensory signals between the GC and the somatosensory cortex reveals that although the somatosensory cortex exhibits higher overall classification accuracy, the GC contains a substantial amount of thermosensory information.

Overall, our results offer a comprehensive analysis of the gustatory cortex’s ability to encode thermal information in addition to gustatory stimuli, highlighting its crucial role in processing thermal cues relevant to taste. This dual capacity emphasizes the importance of incorporating temperature as a key factor in future studies of cortical taste coding.

## Methods

### Experimental design and statistical analysis

#### Data acquisition

The experiments in this study were conducted on 30 wild-type C57BL/6J adult mice (aged 10–20 weeks; comprising 16 males and 14 females). The mice were acquired from The Jackson Laboratory and upon arrival housed in conditions with a 12/12 hour light/dark cycle, with unrestricted access to food and water. The experiments and training sessions were performed during the light phase of this cycle. All experiments are reviewed and approved by the Florida State University Institutional Animal Care and Use Committee (IACUC) under the protocol PROTO202100006. The experimental dataset consists of 962 neurons recorded in the gustatory cortex and 529 neurons recorded in the oral somatosensory cortex. All gustatory cortex (GC) data are from previously published data (Bouaichi et al., 2023; Neese et al., 2022), while recordings from the oral somatosensory cortex are from a new unpublished dataset. For the recording in the gustatory cortex, the taste dataset (529 neurons) neurons come from a previously published dataset (Bouaichi and Vincis, 2020; Neese et al., 2022); neural activity was recorded while the mice were allowed to freely lick to receive 3 *µ*l of one of the four taste stimuli (sucrose, NaCl, citric acid and quinine) at room temperature; the temperature dataset (433 neurons) neurons come from a second previously published dataset (Bouaichi et al., 2023); neural activity was recorded while mice were allowed to freely lick to receive 3 *µ*l of deionized water presented at one of three non-nociceptive temperatures (14°, 25° and 36°C). It is important to note that the experimental settings in which neurons were recorded across the different datasets were identical, with the only difference being the type of stimuli used: taste stimuli for the taste dataset and deionized water at different temperatures for the temperature dataset.

For the recording in the oral somatosensory cortex, mice were anesthetized with an intraperitoneal injection of a cocktail of ketamine (25 mg/ml) and dexmedetomidine (0.25 mg/ml), followed by a subcutaneous injection of Carprofen (20mg/kg). The depth of anesthesia was monitored regularly by visual inspection of breathing rate, whisker reflexes, and tail reflex. Body temperature was maintained at 35°C using a heating pad (DC temperature control system, FHC, Bowdoin, ME).

Once a surgical plane of anesthesia was achieved, the animal’s head was shaved, cleaned, and disinfected with iodine solution and 70% alcohol before positioning it in a stereotaxic plate. A first craniotomy was drilled above the left oral somatosensory cortex on the mouse’s skull (AP: 1.1 mm, ML: 3.8 mm relative to bregma) to implant a movable bundle of eight tetrodes and one single reference wire (Sandvik-Kanthal, PX000004) with a final impedance of 200–300 kΩ for tetrodes and 20–30 kΩ for the reference wire. A second hole was drilled on top of the visual cortex, where a ground wire (A-M Systems, catalog no. 781000) was lowered ∼300 *µ*m below the brain surface. During surgery, the tetrodes and reference wires were lowered 0.2 mm below the cortical surface; they were further lowered ∼200 *µ*m before the first day of recording and ∼80 *µ*m after each recording session. Before implantation, tetrode wires were coated with a lipophilic fluorescent dye (DiI; Sigma-Aldrich), allowing us to visualize the final location of the tetrodes at the end of each experiment. Tetrodes, ground wires, and a head screw (for the purpose of head restraint) were cemented to the skull with dental acrylic. Animals were allowed to recover for a week before the water restriction regimen began and they receive additional daily Carprofen injections (for a total of 3 days; 20mg/kg) from the first postoperative day. The voltage signals from the tetrodes were captured, digitized and filtered by bandpass (300-6000 Hz) using the Plexon OmniPlex system at a sampling rate of 40 kHz. For automated spike sorting, Kilosort by Pachitariu et al. (Pachitariu et al., 2016) was used on a workstation equipped with an NVIDIA GPU, CUDA, and MATLAB. After spike sorting, Phy software facilitated manual curation. Subsequently, the quality metrics and waveform characteristics were determined using scripts based on SpikeInterface (Buccino et al., 2020). Only units exhibiting an overall firing rate greater than 0.3 Hz, a signal-to-noise ratio greater than 3.0 and an ISI violation rate less than 0.2 were considered for further analysis. At the end of the experiment, the mice were deeply anesthetized for the last time and subjected to transcardial perfusion with 30 ml of PBS, followed by 30 ml of 4% paraformaldehyde (PFA). The brains of the subjects were then removed and further fixed in PFA for 24 hours. After fixation, coronal brain sections (100 µm thick) that included the oral somatosensory cortex were prepared using a vibratome (VT1000 S; Leica, Wetzlar, Germany). To highlight the paths taken by the tetrode bundles and probes, brain sections were counterstained with Hoechst 33342 (1:5000 dilution, H3570; Thermo Fisher, Waltham, MA) using standard techniques and then placed on glass slides. The stained sections of the oral somatosensory cortex were examined and imaged with a fluorescence microscope.

For retrograde tracing experiments, mice were anesthetized with an intraperitoneal injection of a cocktail of ketamine (25 mg/ml) and dexmedetomidine (0.25 mg/ml). The depth of anesthesia was monitored regularly by visual inspection of breathing rate, whisker reflexes, and tail reflex. Body temperature was maintained at 35°C using a heating pad (DC temperature control system, FHC, Bowdoin, ME). Once a surgical plane of anesthesia was achieved, the animal’s head was shaved, cleaned, and disinfected with iodine solution and 70% alcohol before positioning it in a stereotaxic plate. Two small craniotomies were drilled above the left GC (+1.1 mm AP; 3.8 mm ML relative to bregma) and the left oral somatosensory cortex (+1.1 mm AP; 3.8 mm ML; relative to bregma). A glass pipette was loaded with cholera toxin subunit B (CTB-488; Thermo Fisher Scientific, Waltham, MA) and lowered into the GC (2.2 mm from the brain surface). We injected 150 nl of CTB-488 at a rate of 2 nl/s using a Nanoject III microinjection pump (Drummond Scientific, Broomall, PA). Following injection, we waited an additional 10 minutes before slowly extracting the glass pipette. A second glass pipette was then loaded with cholera toxin subunit B (CTB-594; Thermo Fisher Scientific, Waltham, MA) and lowered in the oral somatosensory cortex (-0.8 mm from brain surface). We then injected 150 nl of CTB-594 at a rate of 2 nl/s using a Nanoject III as described above and waited an additional 10 minutes before slowly extracting the glass pipette. The scalp of the mouse was then sutured and the animals were allowed to recover from anesthesia. One week after CTB injections, mice were terminally anesthetized and transcardially perfused with 30 ml of PBS followed by 30 ml of 4% paraformaldehyde (PFA). Similarly to what we described earlier for tetrode tracks, the brains were extracted and postfixed with PFA for 24 h, after which coronal brain slices (100-µm thick) were sectioned with a vibratome (VT1000 S; Leica, Wetzlar, Germany). To visualize the anatomical tracers brain slices were counterstained with Hoechst 33342 (1:5000 dilution, H3570; ThermoFisher, Waltham, MA) using standard techniques and mounted on glass slides. Brain sections were viewed and imaged on a fluorescence microscope.

#### Data Pre-Processing and Normalization

Raw spike trains are mathematically represented as 4000-dimensional vectors, with one entry (either 0 or 1) per millisecond of experimental data collection. Lick timings are also represented as vectors of the same dimension with entries of either 0 or 1 reflecting the absence or presence of a lick at each time point. Time 0 is defined as the time of the first lick for which water is present in the spout. Only data after this time point are used in the classification analysis.

The time between two licks, or *lick interval*, is a natural frame of reference to determine the spiking phase. To help in this, we normalize the lick intervals so that all have a *normalized time duration* of 200 time units. Lick intervals that are longer than 200 ms are contracted and those shorter than 200 ms are stretched. This normalization of the lick interval, or time warping, is done uniformly so that the number of spikes per lick interval is not changed and the relative timing of the spikes within a lick interval is maintained. This is illustrated in Fig. 2A,B. The *phase* of the first spike is ≈ 40*/*200, or 0.2. The phase of the last spike is ≈ 170*/*200, or 0.85. In our analysis, we examine the first 5 lick intervals starting at time 0. Therefore, each spike train vector used in the analysis contains 1000 elements: 200 for the time points within each of 5 normalized lick intervals. We refer to each such vector as a *processed spike train vector*.

### Creating Empirical Rate and Phase Distributions

From the processed spike train vectors, we construct approximate phase and rate probability distributions for each neuron that responds to each stimulus. A phase distribution describes the timing of spikes relative to licks in the processed spike train. To construct this, we first form the union as a multiset of the processed spike trains for that neuron/stimulus pair to create one 1000-dimensional vector for each pair. We convert this distribution, defined only at discrete time points, to a continuous distribution. We then remove the rate information by normalizing the distribution to sum to 1, regardless of the number of spikes considered. To construct a rate distribution, we form a histogram (with bin size of 1) of the number of spikes produced per trial.

Transformation to a continuous distribution is done using a Gaussian Kernel Density Estimator (Pedregosa et al., 2011) as follows. The continuous function that interpolates our recordings for a particular neuron/stimuli pair is

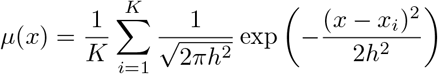

where *x* ∈ [0, 999], and (*x*_1_, *x*_2_, …, *x*_*K*_) are the *K* data points. For phase distributions, the data are the union of all spike times. For rate distributions, the data are the values of the spike frequency histogram. The parameter *h* is called the bandwidth and is akin to a smoothing parameter - the larger the bandwidth, the greater the smoothing. We use *h* = 5 when calculating Phase PDFs and *h* = 2 when calculating Rate PDFs, using a finer bandwidth when smoothing histograms with a small bin size and a larger bandwith when trying to get a sense of the density of spiking for phase distributions. We verified that the results were stable under a range of bandwidth values for phase functions, so we chose a value on the lower end of the stable range to retain as much variability within the data as possible while maintaining similar results.

This process is illustrated in Fig. 2C, D. Panel C shows the timing of spikes over all 5 lick intervals (black bars). These data are used to construct a probability density function (blue curve) using kernel density estimation. Probability values are low because the time spans all five lick intervals, and the number of data points (spikes), *K*, is large over that long time span. Panel D illustrates the construction of the spike-rate density function. The process begins with a histogram of the number of spikes per trial for a particular neuron/stimulus pair. These data are then used in the kernel density estimator to generate a smooth, continuous probability density function (blue curve). In this case, the number of data points, *K*, is the number of different values of spikes per trial that occur in the trials examined.

### Bayesian Classifier Design

We quantify the degree to which a neuron successfully distinguishes one stimulus from the others through classification scores. We employ a Bayesian classification scheme that makes use of one or both of the probability density functions described above. The data sets (processed spike trains) were first divided into training (80%) and testing (20%) sets. The spike trains in the training set were used to construct empirical rate and phase distributions as described above. The spike trains in the test set were used to determine the classification score, as described below.

For classification based on phase distributions, each processed spike train with *N* spikes in the test set is interpreted as *N* i.i.d. samples (*x*_1_, …, *x*_*N*_) from a phase distribution for one of the stimuli *M* stimuli: 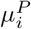, *i* = 1 … *M*. To determine the most likely stimulus (a taste or temperature) 𝒯_*i*_, we maximize *P* (𝒯_*i*_|(*x*_1_, …, *x*_*N*_)) for *i* = 1, …, *M*. Employing Bayes’ theorem, we obtain

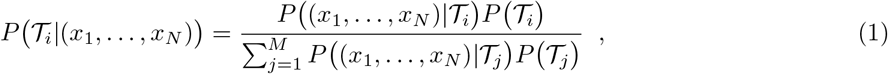

where *P*(𝒯)_*j*_ are the prior probabilities that the observation is from stimulus 𝒯_*j*_. To ensure that all stimuli are equally probable, we include an equal number of trials per stimulus (*Q*) in the analysis, where *Q* is the smallest number of trials recorded for any stimulus in a given data set. For stimuli for which more than *Q* trials were performed, excesses were not included in the analysis. Hence, since all stimuli are equally probable, 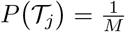. Under the i.i.d. assumption, Eq. (1) simplifies to

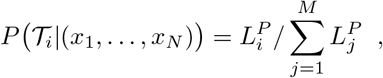

where 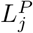 is the likelihood,

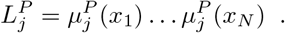

A similar process is used in the case in which rate distributions alone are used in the classification. In this case, each processed spike train is interpreted as a single sample *y* from a rate distribution for one of the stimuli, 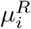, *i* = 1 … *M*. To determine the most likely stimulus 𝒯_*i*_, we maximize *P* (𝒯_*i*_|*y*) for *i* = 1, …, *M*, and from Bayes’ theorem,

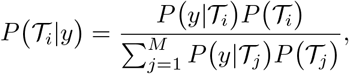

or with the i.i.d. assumption,

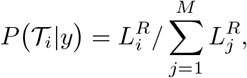

where 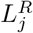 is the likelihood,

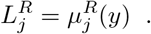

Finally, in the case in which both rate and phase are used in the classification, we maximize the weighted sum of the two probabilities:

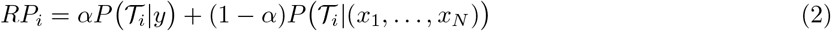

with weight *α* ∈ [0, 1]. When *α* = 0 only the phase information is used and when *α* = 1 only the rate information is used. The values of *α* between 0 and 1 use a combination of the two, and are equally weighted when *α* = 0.5.

Using this approach, we obtain a value *RP*_*i*_ for each of the stimuli, and the stimulus with the highest value is selected as the one most likely to produce the spike train. This is done for all elements of the test set, and the *classification score* is the fraction of times the classification is correct. This process is repeated 15 times for each neuron, each with a different partition of spike trains into training and testing sets. The classification scores are then averaged over these 15 splits to obtain an overall classification score for the neuron.

### Support Vector Machine Classification

To validate our Bayesian analysis classification results, we compare them with a similar classification experiment carried out with a Support Vector Machine (SVM), a tool of classical machine learning (Fan et al., 2008; Pedregosa et al., 2011). Roughly, the SVM determines the optimal hyperplanes that separate the data into a specified number of classes. Each of these is associated with a stimulus, based on the labels of the data used to train the SVM. To perform this experiment, the processed spike train vectors for every neuron/stimulus pair were smoothed by convolving the sequence of values 0 and 1 with a Gaussian kernel (a smoothing window of 100 was used). As in Bayesian classification, the processed spike train data were divided into a training set (80%) and a test set (20%). The training set was used to fit the parameters of the SVM model. Using the trained model, each element of the test set was classified. The fraction of correct classifications is then the classification score. This sequence was repeated 15 times for each neuron, with the same random partitions for training and testing sets that were used in Bayesian classification. The average classification score for a single neuron in all 15 iterations is the final classification score for that neuron. See (Neese et al., 2022) for more details on the SVM classifier and smoothing process.

## Results

### Using a Bayesian classifier to elucidate gustatory cortex activity

To begin investigating how neurons in the gustatory portion of the insular cortex encode oral information in freely licking mice, we made extracellular recordings using movable bundles of tetrodes or silicon probes implanted unilaterally in the GC (Fig. 1). After habituation to head restraint, a group of mildly (1.5 ml per day) water-deprived mice was engaged in a task in which they had to lick a dry spout 6 times to obtain a drop of one of four gustatory stimuli (100 mM sucrose, 50 mM NaCl, 10 mM citric acid, 0.5 mM quinine) and water presented at room temperature. A second group of mice, in a different session, was also trained to receive deionized water at three different non-nociceptive temperatures (14°, 25° and 36°C). At the end of the training (1 week), the recording session started. For the mice included in the taste dataset (n = 12 mice), we recorded and analyzed neural activity (n = 529 neurons) only in response to gustatory stimuli at room temperature (Fig. 1A-B) as discussed in Bouaichi and Vincis (2020) and Neese et al. (2022). For the mice included in the temperature dataset (n = 18 mice), we initially recorded neural activity evoked by taste stimuli presented at room temperature. Following this recording session, we analyze the spiking activity of each neuron. If taste-selective neurons were detected, the tetrodes or probes were lowered approximately 100 *µ*m and subsequent recording sessions exclusively captured the activity evoked by deionized water at various temperatures. As explained in Bouaichi et al. (Bouaichi et al., 2023), this approach was specifically selected to provide additional functional evidence confirming that neuron responses to thermal stimuli were acquired from the taste cortex. Overall, for the temperature dataset we recorded and analyzed 433 GC neurons (Fig. 1A-B; see Methods for additional details).

**Figure 1.**
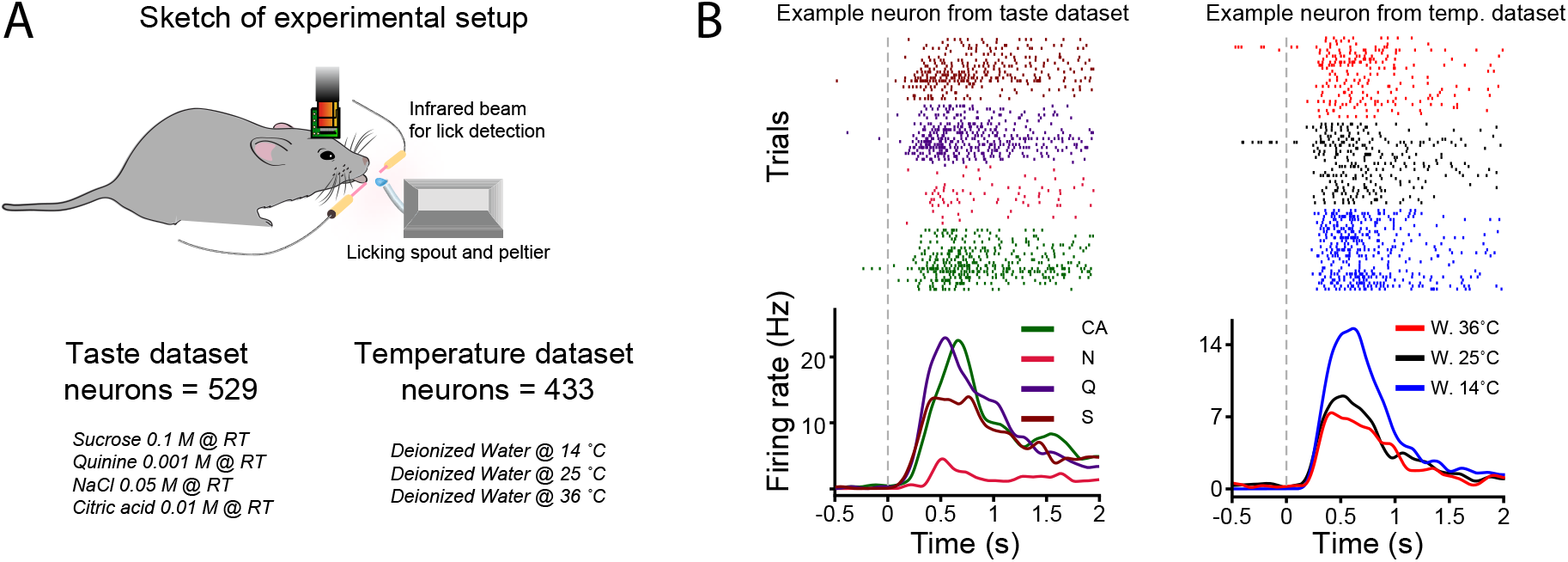
**(A)** Schematic showing the recording setup and a head-restrained mouse licking a spout to obtain oral stimuli. The taste dataset includes GC neurons recorded while animals experience one of four taste stimuli (sucrose 0.1 M, NaCl 0.05 M, citric acid 0.01 M, and quinine 0.001 M) at room temperature. The temperature dataset includes GC neurons recorded while the animals experience deionized water at one of three non-nociceptive temperatures (14°, 25,° 36°C). **(B)** Raster plots and peristimulus time histograms (PSTHs) of 2 representative GC neurons from the taste (left) and temperature (right) dataset.

To investigate the neural dynamics of GC neurons in relation to taste and thermal information, we employed a Bayesian classification approach. For each neuron, the spike trains were initially divided into a training set (80% of total trials for each stimulus) and a test set (20% of total trials for each stimulus). The spike trains from the training sets were then used to construct a spike timing and a spike rate probability distribution function for each neuron-stimulus pair (Fig. 2), which characterize the response of the neuron to the different oral stimuli. For each spike train in the test set, we determined (1) the phase in which each spike occurred and (2) the spike rate for the entire spike train to calculate the corresponding probability values. Then these probability values were combined, weighted by a factor *α* ∈ [0, 1] where *α* < 0.5 places greater emphasis on phase information and *α* > 0.5 gives more weight to rate information. The result is a value *RP*_*j*_, which quantifies the degree to which the spike train in the test set matches the characteristic phase and rate distributions of stimulus *j* in that neuron (Eq. (2)). The neural response is classified as selective to stimulus *j* if the objective function *RP*_*j*_ is greater than that of all other stimuli. In other words, the classification is determined by *RP*_*j*_ > *RP*_*i*_ for all *i* ≠ *j* and *i* = 1, …, *M*, where *M* is the number of different stimuli for that experimental session. The fraction of correctly classified elements in the test set gives the *classification score* for that neuron. We showed previously that a different approach, employing Support Vector Machines (SVM), is highly effective in correctly classifying spiking responses of neurons to a variety of oral stimuli (Neese et al., 2022) in behaving rodents. However, unlike the Bayesian approach we used in this study, the SVM method provides limited information on which specific aspects of the spike trains contribute to the classification success. In contrast, the Bayesian approach employed here not only offers empirical distributions for spiking phases and rates, but also allows for the assignment of different weights to these factors during classification, thereby offering a more detailed understanding of the underlying neural coding mechanisms.

**Figure 2.**
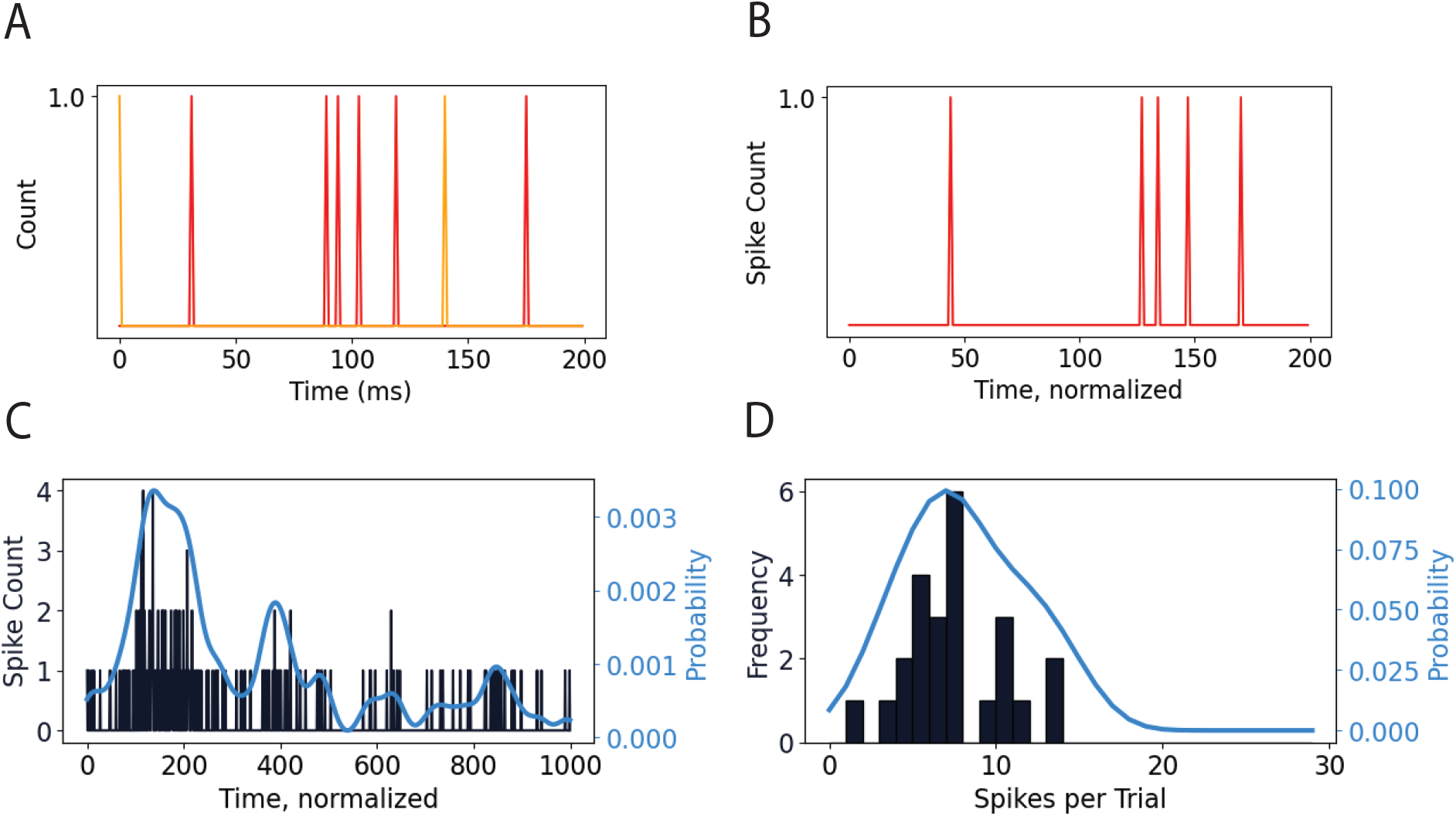
Illustration of the lick interval normalization process and calculation of empirical probability distribution functions. **(A)** A depiction of the timings of the licks (orange) and the neuronal spikes (red). There are 5 spikes within the 140 ms lick interval delimited by the two licks shown. **(B)** The lick interval is extended to 200 ms, preserving the number of spikes and the relative differences in the timing of the spikes. **(C)** For the phase distribution, the black bars indicate the union of all spike times in the processed spike train vectors across all trials. The blue curve is the probability density function generated by kernel density estimation. **(D)** For the rate distribution, the black bars are a histogram showing the frequency of spike numbers per test in a population of processed spike train vectors. The probability density function is again generated using kernel density estimation. This example features data from the 22 trials recorded from neuron 380 in response to sucrose.

Although the proposed Bayesian classification approach provides better insight into the contributions of spike timing and rate, we recognized the need to validate its effectiveness. Specifically, for the sake of comparison, we first compared our new Bayesian method with the SVM approach used in the past (Neese et al., 2022) to ensure its reliability in classifying taste and thermal neural responses. Figure 3 illustrates the comparison of classification scores between the Bayesian and SVM approaches on the same datasets. For each GC neuron, the training sets used to form the empirical rate and phase distributions for the Bayesian analysis were also used to determine the SVM parameters. The test sets were then used to calculate the classification scores for both methods. For the Bayesian classification, “optimal” values of *α*=0.875 and 0.750 were used for the taste (Fig. 3A) and temperature (Fig. 3B) datasets, respectively, with the rationale for these choices discussed in the next section. The classification scores for both datasets indicates that both methods yield comparable scores. Consequently, all subsequent analyzes were performed using the Bayesian classification approach.

**Figure 3.**
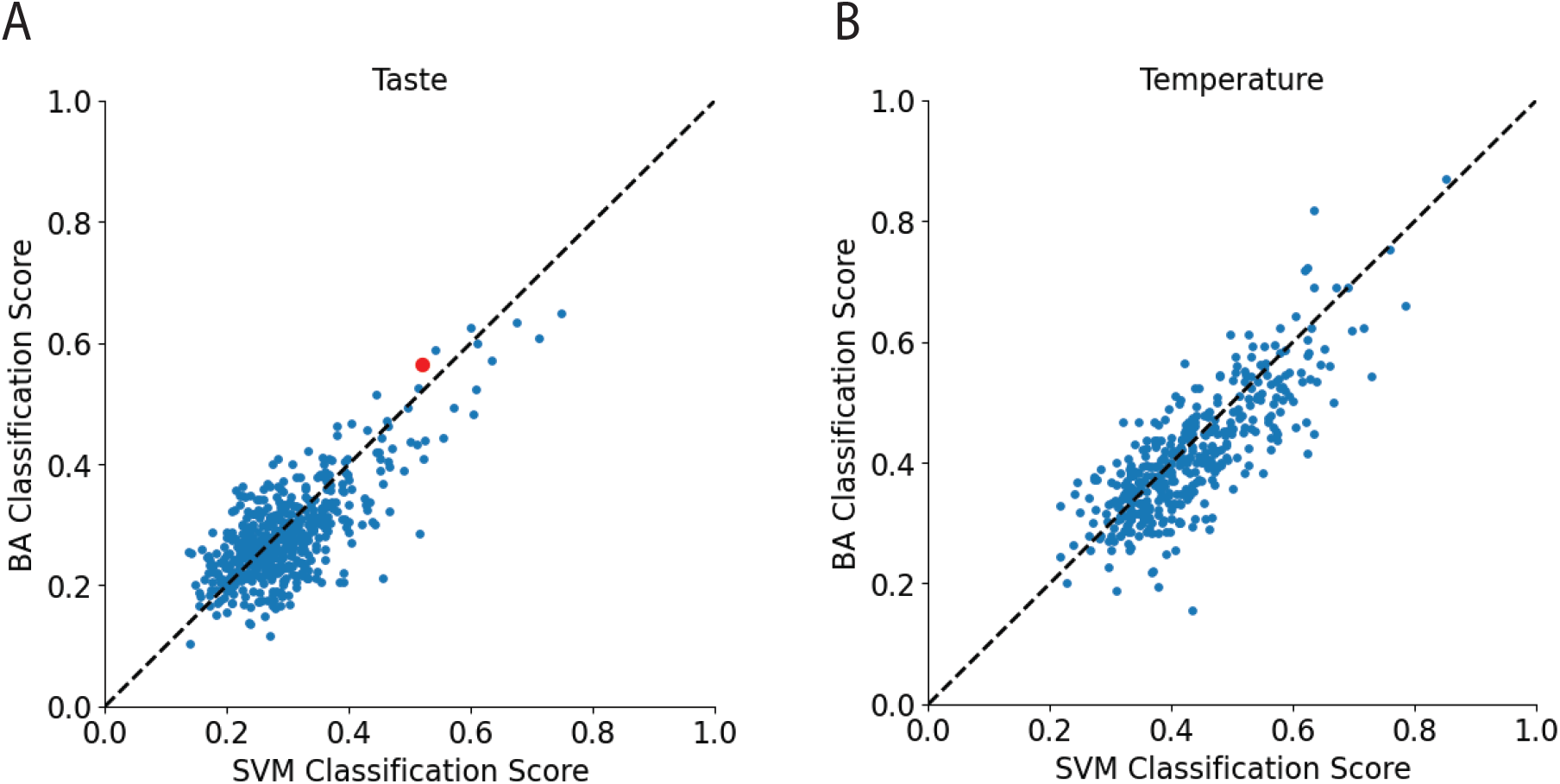
Comparison of classification scores computed with Bayesian and SVM methods. **(A)** The population of 529 neurons in response to 4 tastants (citric acid, NaCl, quinine, and sucrose). The score for random guessing is 0.25. The red point indicates the neuron used in Fig. 2. Points on the dashed line have the same classification score when computed with either method. **(B)** A different population of 433 neurons in response to water at three different temperatures (14°, 25°, and 36°C). The score for random guessing is 0.33. Bayesian analysis scores were computed with *α* = 0.875 (panel A) and *α* = 0.750 (panel B).

### Classification using rate and phase is better than either alone

We first applied the Bayesian classifier to calculate the classification scores for taste information in GC neurons. Classification scores were calculated with different values of *α* ranging from 0 to 1 in increments of 0.125. Previous electrophysiological data obtained in behaving rodents indicate that the fraction of taste-selective neurons in the GC varies depending on multiple factors, including the type of stimulus delivery (intraoral cannulae or active licking), the concentrations of the taste stimulus, the cortical layers, and the method of analysis of stimulus-evoked spike trains (Dikecligil et al., 2020; Levitan et al., 2019; Bouaichi and Vincis, 2020; Katz et al., 2002; Jezzini et al., 2013). In the context of our dataset and experimental conditions, previous analyses have indicated that 10 to 30% of GC neurons can be considered taste-selective (Bouaichi and Vincis, 2020; Neese et al., 2022). Therefore, we focused on the “best” GC neurons, specifically those with taste classification scores in the top 20% (105 neurons). When averaging over this subset of neurons, the mean classification scores ranged from 0.37 to 0.40, significantly higher than the expected score for random guessing (Fig. 4A). The mean score for the classification based solely on phase (*α* = 0) was the lowest among all *α* values. However, although the mean score for the classification based solely on rate (*α* = 1) was higher, it did not reach the maximum value, which was achieved when both rate and phase were combined in the classification (*α* = 0.875). These results confirm that spike timing is a critical factor in taste decoding (Neese et al., 2022) and that gustatory information is enhanced when changes in spike rate are considered in conjunction with temporal dynamics (Katz et al., 2001). Therefore, we used this optimal *α* in all subsequent taste classification analyses unless otherwise noted.

**Figure 4.**
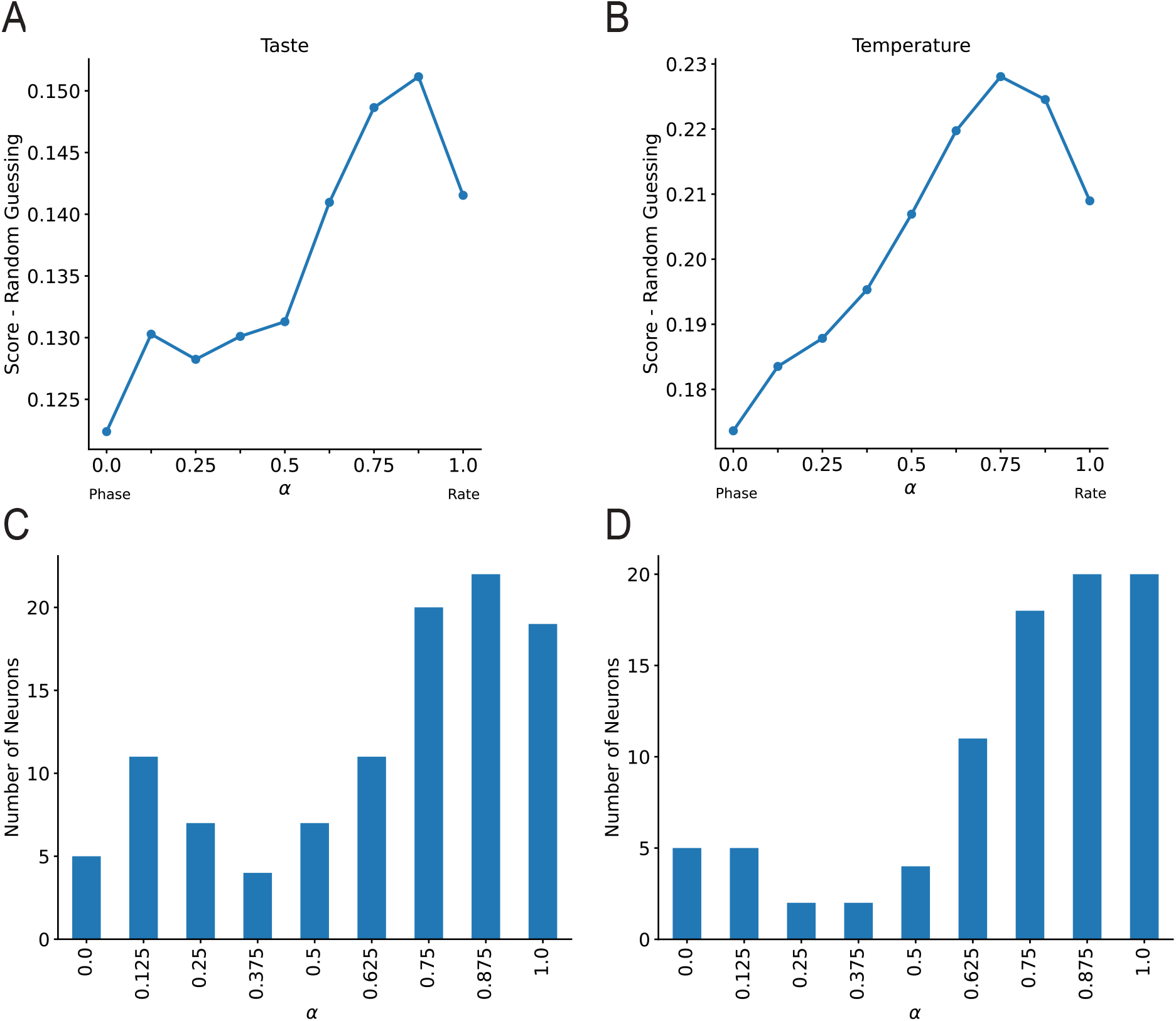
Bayesian-based classification that uses both rate and phase information is optimal. **(A)** Difference between the mean taste classification scores and the random guess score (0.25). For each *α*, only the top 20% neurons were used in the mean classification score. The optimal *α* for this subpopulation is *α* = 0.875. **(B)** Difference between mean classification scores in animals responding to water at three different temperatures (14°, 25°, and 36°C) and random guessing (0.33). The analysis performed was similar to that in panel A. The optimal weighting parameter value for the top 20% subpopulation of neurons is *α* = 0.750. Panels **(C)** and **(D)** show the number of neurons whose highest classification value appears in the top 20% (n=105 for taste, panel **(C)** and 86 for temperature, panel **(D)**) sorted into bins corresponding to which *α* value yields their overall highest classification value.

Next, we aimed to quantify the amount of oral thermal information encoded in the GC spike trains. To do this, we applied the Bayesian classifier to calculate classification scores for fluid temperature using our temperature dataset. For a population of 433 neurons, the mean classification scores ranged from 0.38 to 0.41, slightly above the score of 0.33 expected for random guessing (data not shown). When considering only “best” GC neurons, specifically those with taste classification scores in the top 20% (86 neurons), the mean classification scores were substantially higher, ranging from 0.18 to 0.23 above the expected score for random guessing (Fig. 4B). As in the taste data set, the classification score based on rate alone exceeded that based on phase alone, and the optimal combination of the two involved primarily rate, with phase also contributing to the objective function (*α* = 0.750). This optimal value *α* was used in all subsequent temperature classification analyses.

While Fig. 4A, B demonstrate that the optimal *α* for the neural populations is near 0.75, indicating that putting more weight on spike rate than spike phase produces better classification, it may be that many individual neurons in the population perform best at lower *α* values. To investigate this, for each neuron we performed the classification over the full range of *α* ∈ [0, 1] and picked the *α* that gave the best score. Then out of this neural population we subsampled, considering now only the neurons that scored in the top 20 percentile. These subsampled neurons were then used to construct histograms of the optimal *α* values. The histogram in Fig. 4C corresponds to taste classification with four tastants and shows a peak at *α* = 0.125 and a larger peak at *α* = 0.875. Similarly, for the classification of three water temperatures there was a lower peak near *α* = 0 and a larger peak near *α* = 1. In both cases, then, some neurons perform best using primarily phase, with some contribution from rate, while a larger fraction perform best using primarily rate, with some contribution from phase.

### Many GC neurons are highly responsive to temperature and taste

To go beyond average classification scores, we assessed whether the spike activity of recorded GC neurons decodes chemosensory and thermal stimuli uniformly or whether it more readily distinguishes certain tastes or temperatures. To investigate this, we focused on the “best” GC neurons which are, as explained earlier, those with taste and temperature classification scores in the top 20%.

Figure 5A-B shows the results of this analysis. Each bar’s color coding illustrates how a neuron’s spike train was classified (i.e., predicted stimulus; y-axis), with the x-axis labels indicating the actual stimulus for the spike train. In panel A, for example, more spike trains associated with the citric acid (C) stimulus were correctly classified as citric acid (blue region at the bottom of the first bar) than any other stimulus. When misclassification occurred, the error rate was similar in the other three tastants. Similar patterns were observed for NaCl (N) and quinine (Q). Sucrose (S) had the lowest percentage of correct classifications; the fraction of spike trains incorrectly classified as citric acid was nearly equal to those correctly classified as sucrose. This suggests that the top-performing GC neurons examined were better at distinguishing citric acid, NaCl, and quinine compared to sucrose. A similar analysis was conducted for the temperature dataset (Fig. 5B). Spike trains in response to deionized water at 14°C were correctly classified almost 60% of the time (first bar). The most frequent misclassification for 14°C was at 25°C (yellow region of the first bar). The water at 25°C produced the lowest percentage of correct classifications (second bar), with the majority of misclassifications identifying the cool temperature (14°C). Water at 36°C resulted in the highest percentage of correct classifications (third bar), with the most common misclassification also being water at 25°C (yellow region of the last bar).

**Figure 5.**
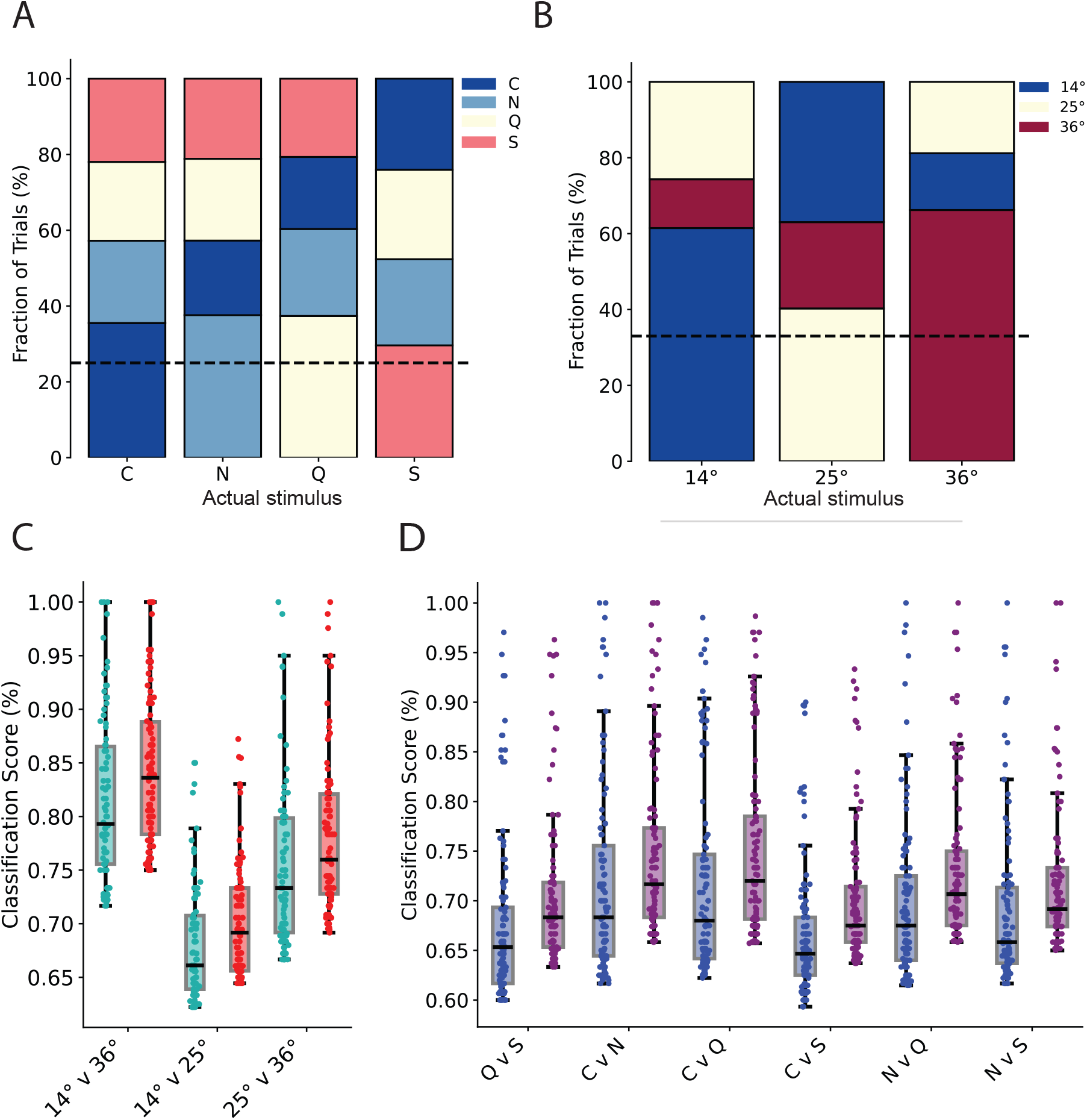
**(A)** Spike train classification from the neurons that performed at the top 20% for taste in the GC (n=105). The color corresponds to the way the spike train was classified. The label on the x-axis is the actual stimulus for the spike train. C=citric acid, N=NaCl, Q=quinine, and S=sucrose. The correct classifications are at the bottom of each bar. The black horizontal dashed line is the expected score for random guessing. **(B)** Spike train classification from the neurons that performed at the top 20% for temperature in the GC (n=86). The fraction of correct classifications was much higher for 14°C and 36° temperatures than for 25°C. **(C)** Comparison of classification scores for pairs of temperatures, using neurons scoring in the top 20%. Teal shows results when selecting *α* = 0.75 for the whole population then subsampling the top 20%, while Red shows results using neuron-specific optimal *α* values and selecting the best-performing 20% of the neurons. **(D)** Comparison of classification scores for pairs of tastes, using only neurons that performed at the top 20%. C=citric acid, N=NaCl, Q=quinine, and S=sucrose. One-way ANOVA between taste pairs: F=6.12, p *<* 0.001. Blue indicates results when selecting *α* = 0.875 for the whole population then subsampling the top 20%, where purple indicates results using the neuron-specific optimal *α* values and selecting the best-performing 20%. For both C and D, the box plots represent the data from Q1 to Q3. The center line in each box indicates the median score, whiskers extend to ±**1.5IQR** and outliers are represented by points outside the range of the whiskers. Overlaid points show the classification score for every neuron considered in the subset.

Next, we compared the distribution of classification scores for pairs of stimuli (Fig. 5C). Visual inspection of the box plots indicates that the mean classification scores obtained for thermal stimuli at 14°C and 36°C, the two thermal stimuli with the highest temperature contrast, are above any other, just below 80%. One-way analysis of variance (ANOVA) tests revealed that the classification scores were significantly different (F = 76.88, p < 0.001). Post hoc pairwise comparison (using the Bonferroni correction method to adjust for multiple tests) further revealed significant differences between the three temperature pairs tested, confirming that each pair exhibited distinct classification scores (Table 1). Classification scores were improved by using the neuron-specific *α* values (red points in Fig. 5C), rather than using a single *α* optimized over the population (teal points), but the improvement is small. The analysis suggests that GC neurons can effectively differentiate between thermal stimuli, excelling in distinguishing both large contrasts (14°C vs. 36°C) and, although less effectively, more subtle temperature differences. A similar analysis, including ANOVA and post hoc tests, was performed on taste stimulus pairs, with the results presented in Fig. 5D and Table 1. Again, there was a small increase in the classification scores when neuron-specific *α* values were used (purple points) rather than a single optimum *α* for the entire population (blue points). Based on a qualitative overview of these data, it could be tempting to speculate that, at least within our datasets, GC neurons seem to encode oral thermal signals more effectively than taste information. However, a quantitative assessment of this point is not appropriate at this time; we will explore this further in the subsequent results sections.

**Table 1:** Table of the p values of pairwise comparisons between oral stimuli using the Bonferroni correction method to adjust for multiple testing (type I error) associated with figure. 5C-D. Significant p values (p *<* 0.05) are indicated in red.

|  | 14°C v 36°C | 14°C v 25°C | Q v S | C v N | C v Q | C v S | N v Q |
| --- | --- | --- | --- | --- | --- | --- | --- |
| 14°C v 25°C | < 0.001 | - |  |  |  |  |  |
| 25°C v 36 °C | < 0.001 | < 0.001 |  |  |  |  |  |
| C v N |  |  | 0.016 | - | - | - | - |
| C v Q |  |  | 0.01679 | 1 | - | - | - |
| C v S |  |  | 1 | < 0.001 | < 0.001 | - | - |
| N v Q |  |  | 1 | 1 | 1 | 0.12378 | - |
| N v S |  |  | 1 | 0.99108 | 1 | 0.52713 | 1 |

### Stimulus identification over individual lick intervals

Thus far, our analysis has focused primarily on spike trains that span five consecutive lick intervals, and the first lick occurs when the stimulus is delivered. However, an intriguing question arises: how robustly are the chemo- and thermosensory stimuli encoded within spike trains over post-stimulus lick intervals compared to the entire five-lick duration? Does classification performance vary significantly between individual lick intervals? To address these questions, we performed a Bayesian classification analysis using portions of processed spike trains that occurred during individual lick intervals (Fig. 6). Similarly to what was described earlier, this analysis used spike trains of “best” neurons for taste or temperature classifications when considering the duration of five licks. Figures 6A and B depict the mean classification score in the different lick intervals for the taste triplets and the temperature, respectively. Visual inspection of the graphs reveals two main points. First, the initial lick interval, which corresponds to the time period approximately 145 ms after stimulus delivery, contains both chemosensory and thermal information. Second, while temperature classification reaches a plateau during the second lick interval, taste classification appears to take longer to reach its peak score, which occurs in the third lick interval. To further investigate this trend, we calculated classification scores for different lick intervals and different rate/phase weighting values *α* (Fig. 6C). For each combination of lick interval and *α*, scores for all neurons were calculated and the top 20% were used to compute the means in this panel. For both taste and temperature, although the optimal *α* varies somewhat from lick interval to lick interval, the performance is best for values of *α* close to, but less than, 1 (Fig. 6C). This indicates that a classification that weighted heavily, but not entirely, on the spike rate is optimal. Furthermore, while the classification scores for temperature beyond the first lick interval are comparable between the *α* values, for taste stimuli the classification score is highest during the third lick interval regardless of *α* (Fig. 6C). Analysis of differences in all classification scores using the second and third lick intervals revealed early success in the classification of thermal stimuli compared to taste stimuli (Fig. 6D; ttest: t(9.4)=3.39, p=0.0073). This suggests different processing of thermal and chemosensory stimuli within the GC, confirming that the timing of neural events can contribute to the representation of oral stimuli (Lemon and Katz, 2007) and that the somatosensory input can be encoded earlier than the chemosensory input (Katz et al., 2001). An important caveat to consider is that the taste stimuli were delivered in solutions at room temperature (around 22°-23°C) and at fixed concentration. Future studies are necessary to investigate whether and to what extent the magnitude and lick-related time course of taste classification vary as a function of temperature and stimulus concentration.

**Figure 6.**
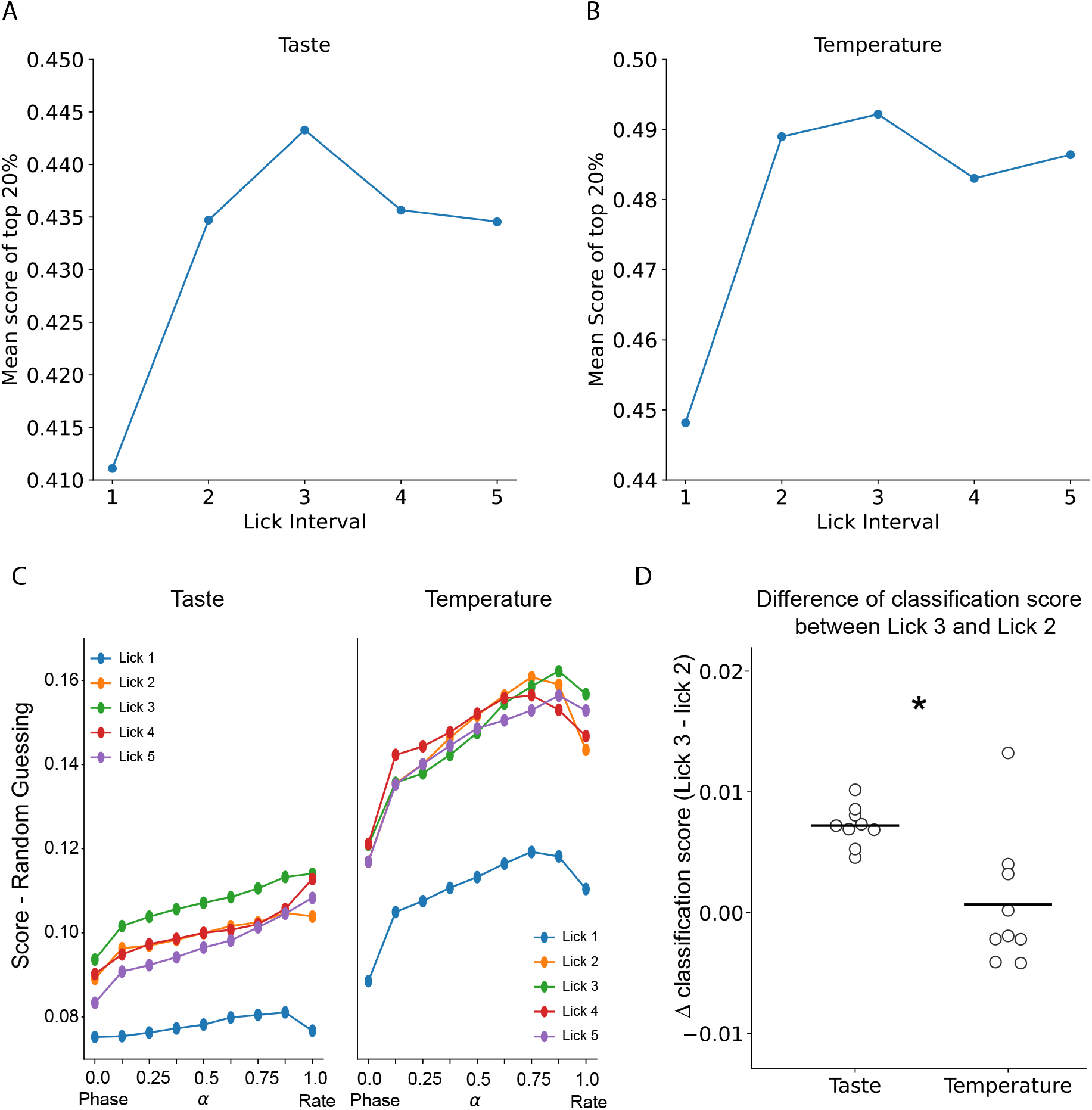
Classification scores based on individual lick intervals. **(A)** Calculated as the average of scores for taste triplets. **(B)** Based on three different water temperatures. Each point represents the mean classification score from the top 20% of neurons when considering classification scores over that particular lick interval only. **(C)** Left panel shows the difference between mean classification score and random guessing for the average of all taste triplets over the 5 different lick intervals across the range of *α* values. The data set from panel A is used in the analysis. The right panel shows a similar analysis, but for the three water temperatures. The data set from panel B is used. **(D)** Difference in classification scores between lick interval 3 and lick interval 2, for taste and thermal stimuli. Each point represents the score difference for a particular value of *α*. The horizontal bars are the average values. *p<0.05.

### Comparison of taste and thermal responses between taste and somatosensory cortex

To summarize our findings so far, we applied a Bayesian analysis to calculate classification scores for GC spike trains, accounting for both the rate and phase codes in response to different oral thermal and gustatory stimuli. Our results appear to indicate that the temperature of the fluid is encoded more effectively by taste cortical neurons, making it more prominent than taste information. However, an important caveat must be acknowledged. Taste responses were recorded exclusively at room temperature (ranging from 22°C to 23°C). Previous work in anesthetized rodents has shown that temperature can have nonlinear additive or subtractive effects on neural activity in response to some tastes (Lemon, 2017), although no information is available for awake behaving animals. Thus, while our results qualitatively suggest that the gustatory cortex reliably encodes oral thermal signals, quantifying this effect compared to taste information is currently not appropriate.

To facilitate a more accurate evaluation of oral thermal coding in the GC, we recorded taste and thermal-evoked spiking activity in oral somatosensory fields as a point of comparison (Fig. 7A). To this end, we implanted a movable bundle of tetrodes unilaterally in the cortical fields dorsal to the GC (Fig. 7B). Anatomical and functional studies confirmed that the cortical area that represents the somatosensory input of the tongue and the intraoral region is located immediately dorsal to the gustatory cortex (Remple et al., 2003; Accolla et al., 2007; Nakamura et al., 2015; Samuelsen and Vincis, 2021; Clemens et al., 2018). Our histological analysis confirmed that the oral somatosensory fields targeted by our tetrodes received thalamic input from the ventroposteromedial (VPM) and not from the gustatory thalamus (VPMpc) (Fig. 7C). Neural recordings with respect to taste and temperature in the somatosensory fields were made under identical conditions used to collect the two data sets in the GC described above (see Methods). A total of 461 neurons from 16 wild-type mice and 68 neurons from 7 mice were recorded for the temperature and taste datasets respectively. Despite the difference in neuron count, the taste dataset serves as an additional physiological control, verifying the location of neurons in the somatosensory cortical fields. Based on previous research, gustatory responses are not expected in this area (Clemens et al., 2018; Accolla et al., 2007). Figure 7D shows the raster plots and PSTHs of three representative neurons. Visual inspection of the plots indicate that, while the neuron in the taste dataset appears to respond similarly to all stimuli (i.e., being modulated by the presence of the fluid irrespective of the taste), the two neurons in the temperature dataset each responded to thermal signals in a selective and temporally dynamic manner. We then used the Bayesian analysis described above to calculate the average taste and temperature classification scores for the neurons recorded in the somatosensory oral fields. Figure 7E shows the plots of the taste (green) and temperature (red) average classification score of the 50 best neurons recorded in the GC and somatosensory cortex (S).

**Figure 7.**
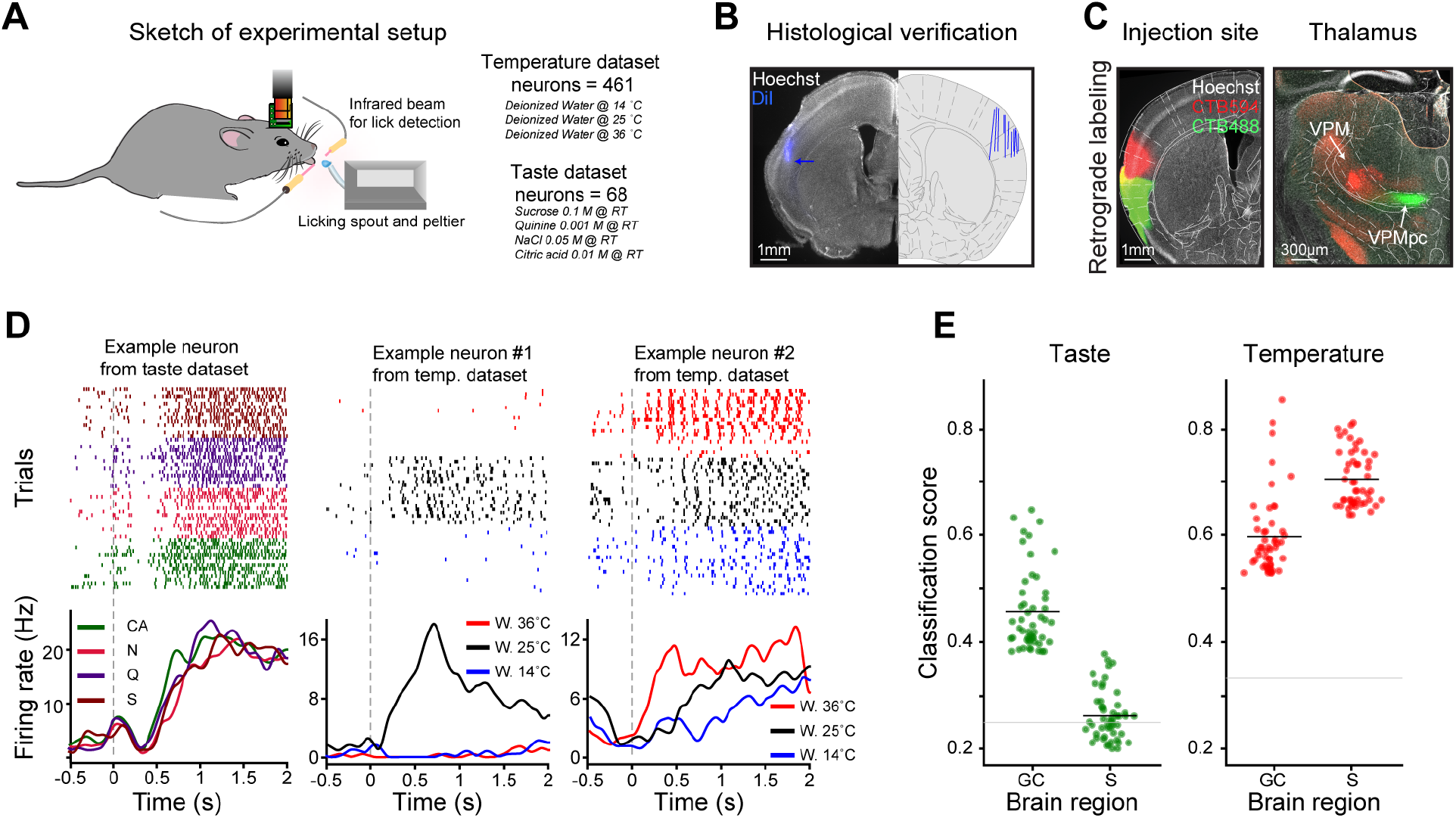
**(A)** Schematic showing the recording setup and a head-restrained mouse licking a spout to obtain different oral stimuli. (**B**) On the left, histological section showing the track (blue) of 1 tetrode bundle in the oral somatosensory field. Blue arrow points to the tip of the tetrode. On the right, schematic of the summary of tetrode tracks from the mice used for the recordings of temperature or taste in the somatosensory cortex. (**C**) On the left, coronal section of mouse brain showing the CTB injection site (in red, CTB594) in the somatosensory cortex and the gustatory cortex (in green, CTB488) counterstained with Hoechst (in gray). On the right, a coronal section showing the location of CTB^+^ neurons in the somatosensory thamalus (VPM, in red) gustatory thalamus (VPMpc, in green). (**D**) Raster plots and peristimulus time histograms (PSTHs) of 3 representative somatosensory neurons from the taste (left) and temperature (center and right) dataset. Trials pertaining to different oral stimuli are grouped together (in the raster plots) and color-coded (both in the raster plots and PSTHs). (**E**) Plots showing the taste (green) and temperature (red) average classification score of 50 best neurons recorded in the GC and somatosensory cortex (S). Black horizontal lines represent the mean while dotted horizontal gray lines represent chance level.

As expected, neurons in the oral somatosensory cortex do not reliably process gustatory information, as their overall classification was not statistically different from random guessing (t-statistic =-1.84, p-value 0.08) and was different from the classification score of the GC neurons (p-value < 2^*−*16^, t-statistic 15.51). Comparison of temperature classification between neurons of the two cortices indicates that the oral somatosensory cortex contains more information about oral thermal stimuli (p-value < 2^*−*16^, t-statistic -8.39), which is expected due to its primary function in processing somatosensory signals from the oral cavity (Accolla et al., 2007; Clemens et al., 2018; Nakamura et al., 2015) (Fig. 7E). Despite the difference, the GC also shows a high classification score for oral thermal information, highlighting its significant role in thermal processing.

Next, we sought to provide a detailed assessment of the classification performance of each thermal stimulus. To do this, we constructed confusion bar charts and analyzed the classification accuracy of neurons in the somatosensory cortical fields for each temperature (Fig. 8A-B). For ease of qualitative comparison, we also re-plotted the bar chart for thermal decoding in the GC in Fig. 5A. The evaluation of the confusion bar chart provides insight into the differences in the average decoding of thermal stimuli between the two cortices, as shown in Fig. 8. A qualitative comparison suggests that the oral somatosensory cortex exhibits greater reliability in distinguishing each absolute temperature, as indicated by the roughly equal fraction of trials correctly classified for each stimulus (bottom section of each bar, panel B). In contrast, as discussed previously, the GC appears to more reliably encode information for the two temperatures with the greatest differences (14°C and 36°C). This observation is also confirmed by the analysis of the classification scores for pairs of thermal stimuli using the best (20%) decoding neurons (86 neurons for the GC and 92 neurons for the somatosensory cortex; Fig. 8C). A two-way analysis of variance (ANOVA) confirmed that there was a significant difference in the classification scores between the two cortices (F-value 151.09, p < 0.001). Similar results are obtained when the best decoding neurons were selected with a different cut-off point (15% and 25%; F-value 113.507, p< 0.001 and F-value 181.91, p< 0.001 respectively). Post hoc pairwise comparisons further confirmed that while both the GC and the oral somatosensory cortex neurons show similar encoding capabilities for some temperature pairs, neurons in the somatosensory cortex consistently performed better at distinguishing all temperature pairs than those in the GC, with the exception of the 25° vs. 36°C pairs (Fig. 8C and Table 2).

**Table 2:**
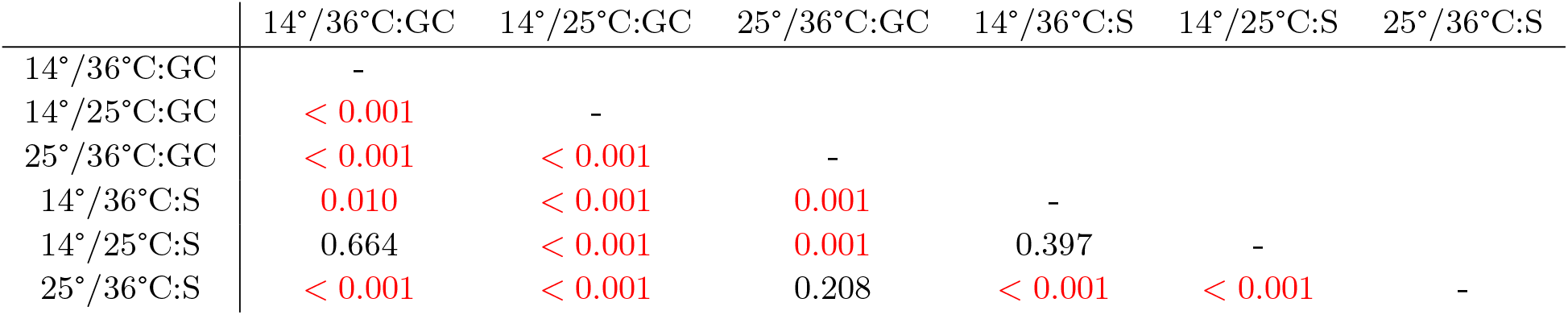
This table presents the p values of post hoc pairwise comparison associated with figure 8C. Significant p values (p < 0.05) are indicated in red.

**Figure 8.**
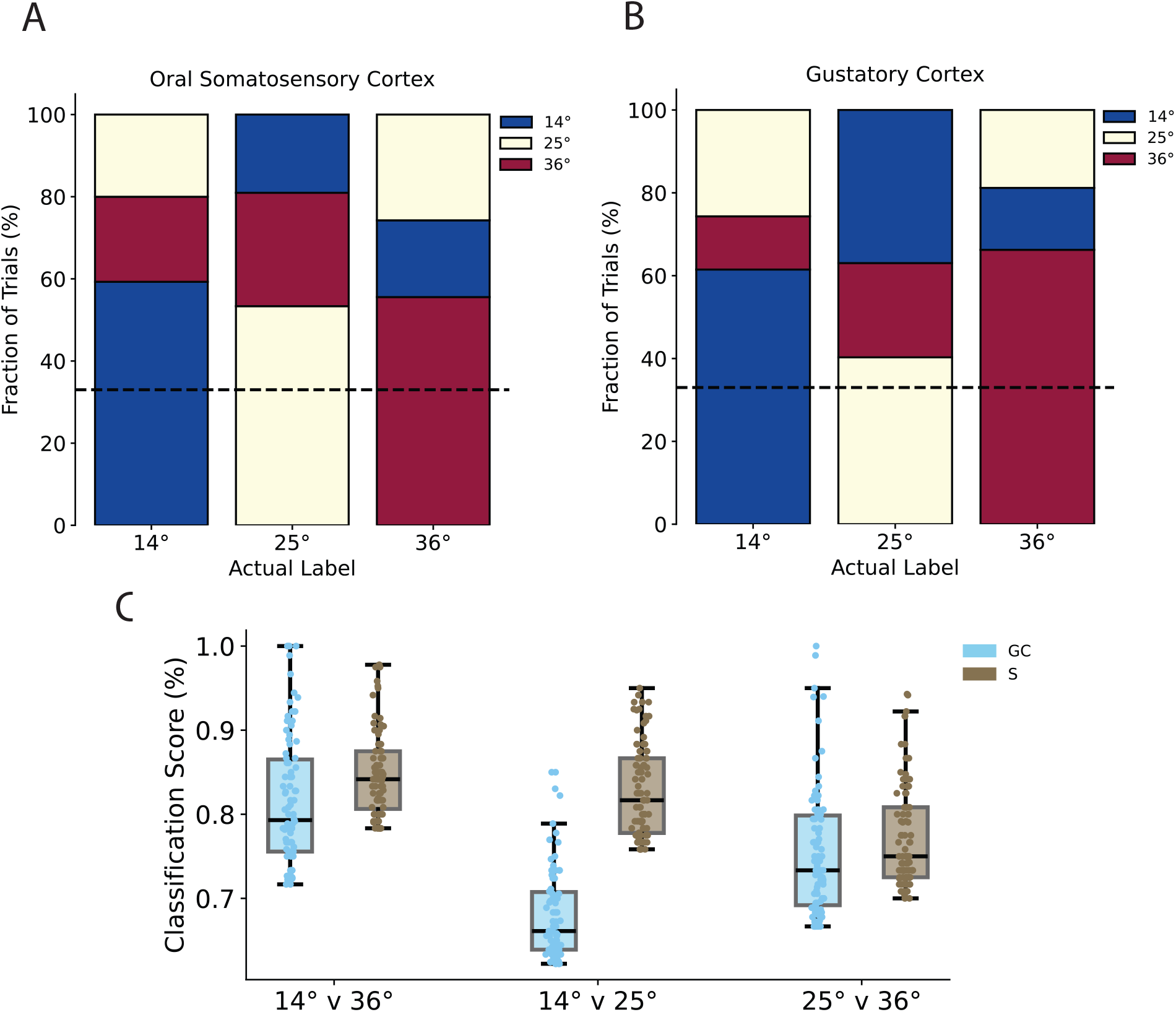
Comparing Thermal Responses **(A)** Spike train classification from the neurons that performed at the top 20% (n = 92)for temperature stimuli in the Oral Somatosensory Cortex. Color of each vertical bar corresponds to the way the spike train was classified. The label on the x-axis is the actual stimulus used to evoke that group of spike trains, and the correct classifications are at the bottom of each bar. The black dashed line represents random guessing (33%). The fraction of correct classifications was much higher for 25°C in the oral somatosensory cortex than in the GC. **(B)** Fig. 5B copied here for ease of qualitative comparison. **(C)** Comparison of classification scores for pairs of temperature stimuli in different brain regions. Box plots represent range of the data from Q1 to Q3, with the median shown as the center line. Whiskers extend to ±1.5*IQR*, with outliers represented as points outside the range of the whiskers. Overlaid points show the classification score for every neuron considered in the subset. GC = gustatory cortex, S = oral somatosensory cortex.

In general, our results indicate that despite the overall higher classification accuracy observed in the somatosensory cortex, the ability of the GC to encode thermal information, particularly for the most distinct temperature pairs, underscores its potential role in integrating thermal cues relevant to gustatory processing.

## Discussion

Experimental evidence from electrophysiological and optical imaging studies shows that neurons in the gustatory cortex represent multimodal, non-gustatory signals experienced before and/or during sampling (Chen et al., 2021; Vincis and Fontanini, 2016; Samuelsen and Fontanini, 2017; Maier, 2017; Livneh et al., 2017; Gardner and Fontanini, 2014; Samuelsen et al., 2012). Of particular relevance are studies that have examined how neurons in the GC respond to intraoral somatosensory characteristics, such as temperature variations. Pioneering work in anesthetized rats (Yamamoto et al., 1981; Kosar et al., 1986) and work in the primate insular/opercular cortex (Verhagen et al., 2004; Kadohisa et al., 2005) indicates that thermal changes in fluid solutions appear to modulate the activity of a subset of GC neurons. Furthermore, our recent work has shown that GC neurons were capable of reliably responding to and discriminating a wide range of innocuous oral temperatures of deionized water in a mostly monotonic manner (Bouaichi et al., 2023). Although these observations highlight the GC’s crucial role in processing thermal signals, they leave open several questions about its capacity to encode oral thermosensory signals and the coding strategies involved. The objective of this study was to evaluate the role of GC neurons in the encoding of oral thermal information and their ability to process chemosensory taste signals at room temperature, particularly compared to the thermosensory and chemosensory coding functions of the oral somatosensory cortex (S), which represents the sensory input of the tongue and intraoral region (Accolla et al., 2007; Nakamura et al., 2015; Clemens et al., 2018; Samuelsen and Vincis, 2021). Given the role of the GC in processing taste stimuli (Mukherjee et al., 2019; Chen et al., 2021; Fletcher et al., 2017; Katz et al., 2001), it could be speculated that taste-related sensory input could inherently have a more prominent representation within this cortical region. This would align with intuitive - but admittedly overly simplistic - expectations, considering the taste cortex function in gustation. However, increasing evidence of nuanced and multifaceted involvement of the GC in the processing of not only gustatory, but also other components of oral stimuli relevant to flavor (Rudenga et al., 2010; Samuelsen and Fontanini, 2017; Verhagen et al., 2004; De Araujo et al., 2003; Vincis and Fontanini, 2016; Bouaichi et al., 2023; De Araujo and Simon, 2009; Maier, 2017) introduces the intriguing possibility that temperature information might play an equally vital role in its sensory integration processes.

In this study, we collected recordings of spiking activity from the gustatory and somatosensory cortex in mice allowed to freely lick to receive a small drop (3 *µ*l) of one of four liquid gustatory stimuli (sucrose, NaCl, citric acid and quinine) at room temperature, or deionized water at one of three different non-nociceptive temperatures (14°, 25° and 36°C). We then developed and employed a new Bayesian-based spike train analysis method to determine the optimal weighting of the rate and phase information for the accurate classification of oral stimuli by cortical neurons. In the gustatory cortex, our analysis revealed that classification scores for both chemosensory and thermosensory modalities were highest when most of the weight was assigned to rate information, although incorporating phase information was necessary to achieve maximum classification accuracy. These results indicate that GC neurons employ a similar strategy to encode oral stimuli across different modalities when experienced via active licking. As we demonstrated previously for taste (Neese et al., 2022), rate information is clearly the dominant factor, but the timing of spikes also plays a complementary role in enhancing thermosensory coding.

Our analysis revealed that the GC robustly encodes thermal information, indicating a strong neural salience of temperature in this brain region. Using the best-performing neurons for encoding (top 20%), our comparisons of classification scores between different thermosensory stimuli revealed that neuronal responses in GC neurons can effectively differentiate between thermal stimuli, excelling in distinguishing both large contrasts (14°C vs. 36°C) and, although less effectively, more subtle temperature differences (Fig. 5C, left panel). A qualitative evaluation of these results might lead to the tempting — though not necessarily valid — deduction that GC neurons encode oral thermal signals differently than taste information. Indeed, Fig. 5 suggests that the GC neurons recorded in this study show a greater capacity to distinguish between deionized water at warm and cool temperatures, particularly at the extremes of our temperature range (14°C vs. 36°C), than between any pair of chemosensory stimuli presented at room temperature. However, this conclusion is highly speculative at this stage for at least two key reasons. First, recent studies show that taste responses in the GC are scattered rather than spatially organized (Chen et al., 2021; Fletcher et al., 2017; Levitan et al., 2019). It is possible that our recordings were mostly from GC regions less sensitive to taste (see, for example, (Chen et al., 2011)). Our experimental approach counters this concern to some extent. Our dataset (Bouaichi and Vincis, 2020; Neese et al., 2022; Bouaichi et al., 2023) includes recordings from a broad section of the GC and captures neurons broadly tuned to different taste qualities (Bouaichi and Vincis, 2020; Neese et al., 2022). However, there may still be variability in the taste responses within our neural sample that could affect the classification outcomes. Second, taste responses were recorded exclusively at room temperature (ranging from 22°C to 23°C) and at fixed concentration. Previous work in anesthetized rodents has shown that temperature and taste concentration can have nonlinear additive or subtractive effects on neural activity in response to some taste qualities (Lemon, 2017). Thus, the outcomes of taste classification could change as a function of the temperature and concentration at which the taste stimulus is presented.

To better evaluate the neural relevance of thermosensory decoding by neurons in the gustatory cortex, we recorded taste and thermal-evoked spiking activity in the somatosensory cortical fields, whose neurons play a crucial role in the encoding and processing of fine somatosensory discrimination (Romo et al., 2002; Foffani et al., 2008; Staiger and Petersen, 2021). In particular, we focused on a region of the somatosensory cortical fields, located immediately dorsally to the gustatory cortex, known to represent the somatosensory input of the tongue, and the intraoral region (Accolla et al., 2007; Nakamura et al., 2015; Samuelsen and Vincis, 2021; Clemens et al., 2018) (Fig. 7). As expected, neurons in the oral somatosensory cortex do not reliably process gustatory information (Clemens et al., 2018). However, in accordance with their primary role in processing somatosensory signals from the oral cavity, they are capable of reliably encoding all different oral temperatures tested. Interestingly, our results revealed a notable difference from findings in a recent study that used calcium imaging to evaluate the cortical representation of thermosensory signals from the skin. In that study, neurons in the somatosensory cortex were reported to respond exclusively to cooling stimuli, with no response to warming stimuli (Vestergaard et al., 2023). In contrast, our study found that all oral thermal stimuli, including the warming temperature of 36°C, are represented and encoded in the somatosensory cortex. The results shown in Fig 8 revealed that although both the gustatory and the somatosensory cortex reliably encode all temperature pairs well above chance level, the somatosensory cortex generally outperforms the GC with the exception of distinguishing 25° vs. 36°C.

Some key questions emerge from our results. First, what functional role could thermal oral signals, encoded by gustatory cortex neurons, play in the context of sensory processing and behavioral responses? We can safely speculate that these oral signals are integral to flavor processing and behavioral responses, particularly in relation to consummatory behavior. The gustatory cortex has been implicated in functions related to taste processing (Katz et al., 2001; Blonde et al., 2015; Sadacca et al., 2012), taste learning (Schier et al., 2016; Kayyal et al., 2021; Yiannakas et al., 2021; Arieli et al., 2022) and expectation (Samuelsen et al., 2012; Gardner and Fontanini, 2014; Vincis and Fontanini, 2016; Livneh et al., 2017), but also in taste-based decision making (Mukherjee et al., 2019; Vincis et al., 2020), highlighting its role not only in sensory processing, but also in modulating consummatory behaviors. Therefore, encoding oral thermal signals by the GC could help improve the fidelity of food evaluation processes, integrating temperature as a crucial parameter in evaluating food quality and palatability (Zellner et al., 1988; Moskowitz, 1973; Torregrossa et al., 2012).

This leads to the second question: why does the gustatory cortex, as opposed to only the somatosensory cortex, appear to play a crucial role in encoding thermal oral signals? Previous experimental evidence may provide information. A study in humans (Craig et al., 2000) and a more recent study in rodents (Vestergaard et al., 2023) indicated that the posterior insular cortex, a region located closely but posterior to the areas targeted in our recordings, functionally represents and underlies the thermosensory perception of the skin. This, together with our findings, points to the possibility of a postero-anterior gradient across the insular cortex for representing thermal information, extending from extraoral to oral thermal signals. This gradient could imply that the gustatory cortex, located within this continuum, is optimally positioned to integrate oral thermal information as a component of its broader role in sensory processing. This specialization might reflect an evolutionary adaptation, prioritizing the gustatory cortex for oral thermal sensing due to its direct implications for feeding and the evaluation of food stimuli, in contrast to the more general body temperature regulation functions possibly attributed to more posterior regions of the insular cortex.

Finally, we consider the potential differences between the gustatory and somatosensory cortices in encoding oral thermal stimuli. One speculative distinction between how the gustatory and somatosensory cortices encode non-nociceptive oral thermal information might lie in their respective roles in processing “gross” versus “fine” temperature differences. It is possible that the gustatory cortex is primarily responsible for encoding broad or “gross” differences in oral thermal stimuli, such as differentiating between temperatures that are cooler or warmer (i.e., 14° vs. 36°C oral stimuli in our experiments) than the “resting” oral temperature. This kind of thermal information, potentially related to GC neurons having a large receptive field for oral thermal stimuli, could be integrated with gustatory signals to influence overall flavor perception based on temperature, potentially enhancing or diminishing certain tastes. However, the GC may be less able to detect fine differences in temperature, where the temperature variations are smaller and closer to the normal oral temperature range. This finer discrimination could instead be the domain of the somatosensory oral cortex, which is well-suited for fine-tuned tactile stimulus discrimination (Romo et al., 2002; Foffani et al., 2008; Staiger and Petersen, 2021), that might lead to including small receptive fields and precise temperature detection. In this context, while the gustatory cortex might primarily handle broad thermal cues to adjust flavor perception, the somatosensory oral cortex could be tuned more to detect and process fine thermal differences, ensuring a more detailed and accurate thermal perception that complements the gustatory experience. Moreover, it is important to consider that these two cortices are likely to interact and should not be viewed as completely independent in their functions. Reciprocal cortico-cortical projections between the GC and the oral somatosensory cortex (Shi and Cassell, 1998) suggest that they work together, likely integrating broad and fine thermal cues to produce a cohesive perception of oral temperature, a critical somatosensory signal, and flavor component. Future studies will be crucial to further elucidate the specific contributions of these two cortices to thermal processing, providing a clearer understanding of how they collaborate to shape sensory perception.

Overall, our findings highlight that GC neurons are not limited to processing taste information; they also play a significant role in encoding thermal signals from the oral cavity. This dual capacity is particularly relevant compared to the encoding observed in the oral somatosensory cortex. These results suggest that temperature should be considered a critical dimension in future studies of cortical taste coding, as it can substantially affect the cortical coding of gustatory stimuli and refine our understanding of the role of the gustatory cortex in shaping nuanced perception of flavor.

## Data availability statement

Data sets generated and/or analyzed during the current study are available in the vincisLab GitHub repository.

